# ETV2 upregulation marks the specification of early cardiomyocytes and endothelial cells during co-differentiation

**DOI:** 10.1101/2022.11.15.516686

**Authors:** Xu Cao, Maria Mircea, Gopala Krishna Yakala, Francijna E. van den Hil, Marcella Brescia, Hailiang Mei, Christine L. Mummery, Stefan Semrau, Valeria V. Orlova

## Abstract

The ability to differentiate human induced pluripotent stem cells (hiPSCs) efficiently into defined cardiac lineages, such as cardiomyocytes and cardiac endothelial cells, is crucial to study human heart development and model cardiovascular diseases *in vitro*. The mechanisms underlying the specification of these cell types during human development are not well-understood which limits fine-tuning and broader application of cardiac model systems. Here, we used the expression of ETV2, a master regulator of hematoendothelial specification in mice, to identify functionally distinct subpopulations during the co-differentiation of endothelial cells and cardiomyocytes from hiPSCs. Targeted analysis of single-cell RNA sequencing data revealed differential ETV2 dynamics in the two lineages. A newly created fluorescent reporter line allowed us to identify early lineage-predisposed states and show that a transient ETV2-high state initiates the specification of endothelial cells. We further demonstrated, unexpectedly, that functional cardiomyocytes can originate from progenitors expressing ETV2 at a low level. Our study thus sheds light on the *in vitro* differentiation dynamics of two important cardiac lineages.

**SIGNIFICANCE STATEMENT:** *In vitro* differentiation of cardiac cell types is of great importance for understanding heart development, disease modeling and future regenerative medicine. Currently, underlying molecular mechanisms are incompletely understood, which limits the efficiency and fine-tuning of present differentiation protocols. Here, we investigated the master regulator ETV2 and showed that its upregulation marks the specification of two cardiac cell types during co-differentiation. Using single-cell RNA-seq and a new fluorescent reporter line we identified lineage-predisposed subpopulations in the ETV2+ cells. We thus resolved ETV2 dynamics at the single-cell level in the context of *in vitro* human cardiac differentiation.

## INTRODUCTION

*In vivo*, cardiomyocytes (CMs) and endothelial cells (ECs) originate from *Mesp1*+ progenitors specified during gastrulation. In mice, these cells appear in the primitive streak and subsequently migrate towards the lateral plate mesoderm around E6.5 [1–4]. The timing of segregation of CMs and ECs from their common progenitor is still controversial. Single-cell RNA-seq (scRNA-seq) of mouse *Mesp1*+ progenitors collected at E6.75 and E7.25 showed that these cells were already segregated into distinct cardiovascular lineages, including CMs and ECs[5]. However, other studies showed that multipotential progenitors were still present in *Flk-1*-expressing lateral plate mesoderm [6,7]. These cells were the first to be recognized as multipotent cardiac progenitor cells (CPCs) [8]. Studies in mouse and chick showed that CPCs come from two different sources [9,10]: the first- and the second heart field (FHF, SHF). The FHF in the cardiac crescent contributes to the primitive heart tube, which serves as a scaffold into which SHF cells can migrate before heart chamber morphogenesis. It has been shown that cells from the SHF are patterned before migration to give rise to different parts of the heart [3,11]. CPCs from FHF and SHF can be distinguished by the expression of ISL1, which is specific to the SHF [12]. *Nkx2-5*-expressing CPCs in both FHF and SHF from E7.5 to E7.75 contribute to both CMs and ECs in the heart [13]. ETS Variant Transcription Factor 2 (Etv2) is a master regulator of endothelial and hematopoietic cell lineages during early development [14]. Etv2 functions downstream of BMP, WNT and NOTCH signaling pathways [15] and regulates the expression of early EC-specific markers, such as *Tal1, Gata2, Lmo2, Tek, Notch1, Notch4* and *Cdh5* [15–18]. In mouse embryonic stem cells (ESCs), VEGF-FLK1 signaling upregulates ETV2 expression to induce hemangiogenic specification via an ETV2 threshold-dependent mechanism [19]. ETV2 expression was also found to direct the segregation of hemangioblasts and smooth muscle cells (SMCs) in mouse ESCs [20].

In human heart development, much less is known about the specification of endothelial and myocardial lineages from multipotent CPCs, both in terms of timing and gene regulatory mechanisms. More specifically, it is still unclear whether ETV2 also plays a role in the segregation of ECs and CMs from CPCs in humans. Overexpression of ETV2 converts human fibroblasts into endothelial-like cells [21] and ETV2 expression levels have been modified in several studies to drive hiPSCs towards ECs in 2D and 3D cultures [22–28]. Paik et al. performed scRNA-seq analysis of hiPSC-derived ECs (hiPSC-ECs), which made up less than 10% of the cells that expressed the cardiac maker *TNNT2*. The developmental dynamics of ECs and cardiac lineages as such were not further studied [29]. In an scRNA-seq study of hiPSC-ECs obtained using a different differentiation protocol [30], ECs were collected at multiple time points. This study showed that endothelial and mesenchymal lineages have a common developmental origin in mesoderm cells but the identity and differentiation potential of these cells was not described.

Previously, we found that *MESP1*+ progenitors derived from human ESCs could give rise to CMs, ECs and SMCs [31,32]. We also developed a co-differentiation system for ECs and CMs from hiPSCs through a common cardiac mesoderm precursor [33]. Here we performed scRNA-seq analysis of this co-differentiation system to elucidate the relationship between *ETV2* expression and specification of ECs and CMs from cardiac mesoderm. *ETV2* expression was observed principally as an initial “pulse” in the endothelial lineage but also in a subpopulation of the myocardial lineage. Using a newly generated ETV2^mCherry^ hiPSC reporter line, we purified two subpopulations of ETV2+ cells and revealed their derivative EC and CM expression characteristics by bulk RNA-seq. These sorted populations also showed distinct differentiation potentials towards CMs and ECs upon further differentiation with VEGF. In summary, this study detailed *ETV2* dynamics during segregation of human CMs and ECs differentiated from hiPSCs.

## MATERIAL AND METHODS

### hiPSC culture

The NCRM1 hiPSC line (NIH Center for Regenerative Medicine (NIH CRM), obtained from RUDCR Infinite Biologics at Rutgers University, hPSCreg number CRMi003-A) was used in this study, except for the single-cell RNA-seq, which was done with LUMC0020iCTRL06 (hPSCreg number LUMCi028-A). hiPSC control lines were cultured in TeSR-E8 on Vitronectin XF and routinely passaged once a week using Gentle Cell Dissociation Reagent (all from Stem Cell Technologies). Prior to targeting, NCRM1 hiPSCs were passaged as a bulk on feeders in hESC medium [34]. RevitaCell (Life Technologies) was added to the medium (1:200) after every passage to enhance viability after single cell passaging with TrypLE (Life technologies).

### Generation of hiPSC reporter line using CRISPR/Cas9

The p15a-cm-hETV2-P2A-NLS-mCherry-neo repair template plasmid was generated using overlap PCR and restriction-based cloning and ligation. The ETV2 homology arms were amplified from genomic DNA and the neomycin cassette flanked by two flippase recognition target (FRT) sites was amplified from the P15 backbone vector (kindly provided by Dr. Konstantinos Anastassiadis, Technical University Dresden). P2A-NLS-mCherry double-stranded DNA fragment was ordered from IDT. The sgRNA/Cas9 plasmid was modified from SpCas9-2A-Puro V2.0 plasmid (Addgene, Feng Zang).

NCRM1 hiPSCs were passaged at a 1:2 or 1:3 ratio into 60 mm dishes to reach 60-70% confluence the next day for transfection. 20 μl lipofectamine (Invitrogen), 8 μg of repair template and 8 μg of sgRNA/Cas9 plasmid were diluted in 600 μl of Opti-MEM and added to each 60 mm dish. After 18 hours the medium was changed to hESC medium. After another 6 hours, G-418 (50 μg/ml) selection was started and was continued for 1 week. Surviving cells were cultured in hESC medium and passage into 6-well plates for the transfection of Flp recombinase expression vector to remove the neomycin cassette [35]. 300 μl of Opti-MEM containing 10 μl lipofectamine and 4 μg CAGGs-Flpo-IRES-puro plasmid was added per well for 18 hours. Puromycin (0.5 μg/ml) selection was started 24 hours post transfection for 2 days. Once recovered, cells were passaged into 96-well plates for clonal expansion via limiting dilution. Targeted clones were identified by PCR and sequencing. Primers outside the ETV2 homology arms and primers inside the targeting construct were used to confirm on-target integration. The absence of mutations within the inserted sequence and untargeted allele was confirmed by Sanger sequencing (BaseClear).

### Endothelial and myocardial lineage co-differentiation from hiPSCs

Endothelial and cardiac cells were induced from hiPSCs in monolayer culture using the CMEC protocol described previously [33]. Briefly, hiPSCs were split at a 1:12 ratio and seeded on 6-well plates coated with 75 μg/mL (growth factor reduced) Matrigel (Corning) on day −1. On day 0, cardiac mesoderm was induced by changing TeSR-E8 to BPEL medium [36], supplemented with 20 ng/mL BMP4 (R&D Systems), 20 ng/mL ACTIVIN A (Miltenyi Biotec) and 1.5 μM CHIR99021 (Axon Medchem). On day 3, cells were refreshed with BPEL supplemented with 5 μM XAV939 (Tocris Bioscience) and 50 ng/ml VEGF (R&D Systems). From day 6 onwards, cells were refreshed every 3 days with BPEL medium supplemented with 50 ng/ml VEGF.

### Fluorescence-activated cell sorting

For FACS sorting on day 4, 5, 6 and 8 of the CMEC protocol, CD144+mCherry+ (DP) and CD144-mCherry+ (SP) cells were sorted using a FACSAria III (BD-Biosciences). Around 20k cells/cm^2^ were seeded on fibronectin-(from bovine plasma, 5μg/ml, Sigma Aldrich) coated plates. Cells were cultured in BPEL supplemented with VEGF (50 ng/ml) until day 10. The medium was refreshed every 3 days. For FACS sorting on day 7 of the CMEC protocol, CD144+mCherry+ (DP), CD144-mCherry+ (SP) and CD144-mCherry-(DN) cells were sorted using a FACSAria III. 1 million cells were seeded in each well of Matrigel-coated 12-well plates in BPEL supplemented with VEGF (50 ng/ml) and RevitaCell (1:200). The medium was refreshed 24 h after seeding and every three days afterwards with BPEL supplemented with VEGF (50 ng/ml).

### Immunofluorescence staining and imaging

Cultured cells were fixed in 4% paraformaldehyde for 15 min, permeabilized for 10 min with PBS containing 0.1% Triton-X 100 (Sigma-Aldrich) and blocked for 1h with PBS containing 5% BSA (Sigma-Aldrich). Then cells were stained with primary antibody overnight at 4°C. The next day, cells were washed three times (20 min each time) with PBS. After that, cells were incubated with fluorochrome-conjugated secondary antibodies for 1h at room temperature and washed three times (20 min each time) with PBS. Then, cells were stained with DAPI (Life Technologies) for 10 min at room temperature and washed once with PBS for 10min. Both primary and secondary antibodies were diluted in 5% BSA/PBS. Images were taken with the EVOS FL AUTO2 imaging system (ThermoFischer Scientific) with a 20x objective, or using the Incucyte^®^ system (Sartorius). Confocal imaging was done using a Leica SP8WLL confocal laser-scanning microscope using a 63x objective and z-stack acquisition. Details of all antibodies used are provided in Table S1.

### FACS analysis

Cells were washed once with FACS buffer (PBS containing 0.5% BSA and 2 mM EDTA) and stained with FACS antibodies for 30 min at 4oC. Samples were washed once with FACS buffer and analyzed on the MACSQuant VYB (Miltenyi Biotech) equipped with a violet (405 nm), blue (488 nm) and yellow (561 nm) laser. The results were analyzed using Flowjo v10 (FlowJo, LLC). Details of all fluorochrome conjugated FACS antibodies are provided in Table S1.

### Quantitative Real-Time Polymerase Chain Reaction (qPCR)

Total RNA was extracted using the NucleoSpin^®^RNA kit (Macherey-Nagel) according to the manufacturer’s protocol. cDNA was synthesized using an iScript-cDNA Synthesis kit (Bio-Rad). iTaq Universal SYBR Green Supermixes (Bio-Rad) and the Bio-Rad CFX384 real-time system were used for the PCR reaction and detection. Relative gene expression was determined according to the standard ΔCT calculation and normalized to housekeeping genes (mean of HARP and RPL37A). Details of all primers used are provided in Table S2.

### Bulk RNA sequencing and analysis

Cells were sorted on differentiation day 4, 5, 6 and 8 for bulk RNA-Seq. Total RNA was extracted using the NucleoSpin^®^ RNA kit (Macherey-Nagel). Whole transcriptome data were generated at BGI (Shenzhen, China) using the Illumina Hiseq4000 (100bp paired end reads). Raw data was processed using the LUMC BIOPET Gentrap pipeline (https://github.com/biopet/biopet), which comprises FASTQ preprocessing, alignment and read quantification. Sickle (v1.2) was used to trim low-quality read ends (https://github.com/najoshi/sickle). Cutadapt (v1.1) was used for adapter clipping [37], reads were aligned to the human reference genome GRCh38 using GSNAP (gmap-2014-12-23) [38,39] and gene read quantification with htseq-count (v0.6.1p1) against the Ensembl v87 annotation [40]. Gene length and GC content bias were normalized using the R package cqn (v1.28.1) [41]. Genes were excluded if the number of reads was below 5 reads in ≥90% of the samples. The final dataset consisted of gene expression levels of 31 samples and 22,419 genes.

Differentially expressed genes were identified using generalized linear models as implemented in edgeR (3.24.3) [42]. P-values were adjusted using the Benjamini-Hochberg procedure and FDR ≤ 0.05 was considered significant. Analyses were performed using R (version 3.5.2). The PCA plot was generated with the built-in R function prcomp using the transposed normalized RPKM matrix. Correlation among samples was calculated using the cor function with spearman method and the correlation heatmap was generated with aheatmap function (NMF package).

Gene clusters were calculated with the CancerSubtypes package [43]. The top 3000 most variable genes across all chosen samples were identified based on the Median Absolute Deviation (MAD) using the FSbyMAD function, then expression was standardized for each gene. K clusters were calculated using k-means clustering with Euclidean distance. Clustering was iterated 1000 times for each k in the range of 2 to 10. Heatmaps of genes in all clusters were generated using the base R heatmap function. Gene ontology enrichment for each cluster was performed using the compareCluster function of clusterProfiler package (v3.10.1) [44] and q ≤ 0.05 was considered significant.

### Single-cell RNA sequencing and analysis

#### Sample preparation and sequencing

Cells were dissociated into single cells on day 6 of CMEC differentiation and loaded into the 10X Chromium Controller for library construction using the Single-Cell 3’ Library Kit. Next, indexed cDNA libraries were sequenced on the HiSeq4000 platform. Mean reads per cell were 28,499 in the first replicate and 29,388 in the second replicate.

#### Pre-processing

Both replicates of day 6 CMEC differentiation were merged into one data set. The average number of detected genes was 2643 and the average total expression per cell was 10382 (Figure S1A-B). Then, undetected genes (> 1 UMI count detected in less than two cells) and cells with low number of transcripts were removed from further analysis (Figure S1A-B). This resulted in 5107 cells for the first replicate, and 3743 cells for the second replicate and 13243 genes. Expression profiles were normalized with the R package *scran* (V 1.10.2) using the method described in [45]. The 5% most highly variable genes (HVGs) for each replicate were calculated with scran after excluding ribosomal genes (obtained from the HGNC website without any filtering for minimum gene expression), stress-related genes [46] and mitochondrial genes. For downstream analysis the top 5% HVGs were used after excluding proliferation [47] and cell cycle [48] related genes.

#### Cell cycle analysis

For each data set, cell cycle analysis was performed with the *scran* package using the *cyclone* function [49] on normalized counts. Cells with a G2/M score higher than 0.2 were considered to be in G2/M phase. Otherwise, they were classified as G1/S. Using this binary classifier as predictor, we regressed out cell cycle effects with the R package *limma* (V 3.42.2) [50] applied to log-transformed normalized counts. The two replicates were then batch corrected with the fast mutual nearest neighbors (MNN) correction method [51] on the cell cycle corrected counts, using the 30 first principal components and 20 nearest-neighbors.

#### Clustering

Batch-corrected counts were standardized per gene and then used to create a shared nearest neighbour (SNN) graph with the *scran* R package (d = 30, k = 20). Louvain clustering was applied to the SNN graph using the *igraph* python package (V 0.7.1) with 0.4 as the resolution parameter. This resulted in 5 clusters. Two of these 5 clusters were excluded from further analysis based on the expression of pluripotency markers [50].

#### Dimensionality reduction and pseudotime

Dimensionality reduction was performed using the python *scanpy* pipeline (V 1.4.6). Firstly, a 20 nearest-neighbors (knn, k=20) graph was created from diffusion components of the batch corrected data sets. Diffusion components are the eigenvectors of the diffusion operator which is calculated from Euclidean distances and a Gaussian kernel. The aim is to find a lower dimensional embedding that considers cellular dynamics. The graph was projected into two dimensions with the default force-directed graph layout and starting positions obtained from the partition-based graph abstraction (PAGA) algorithm [52]. PAGA estimates connectivities between partitions and performs an improved version of diffusion pseudotime. Diffusion pseudotime [51,52] was calculated on these graphs with root cells selected from the “Cardiac Mesoderm” cluster.

Average gene expression trajectories were calculated by dividing the cells of each cluster into bins along pseudotime. 50 bins were created for cardiac mesoderm and 30 bins each for ECs and CMs. The average log-expression per bin was then calculated. The value of the threshold indicated in Fig. 1 D,E was determined by calculating the point in pseudotime where the average ETV2 expression was the lowest in the endothelial cell cluster before the peak expression, which corresponds to a value around 0.25.

**Figure 1.**
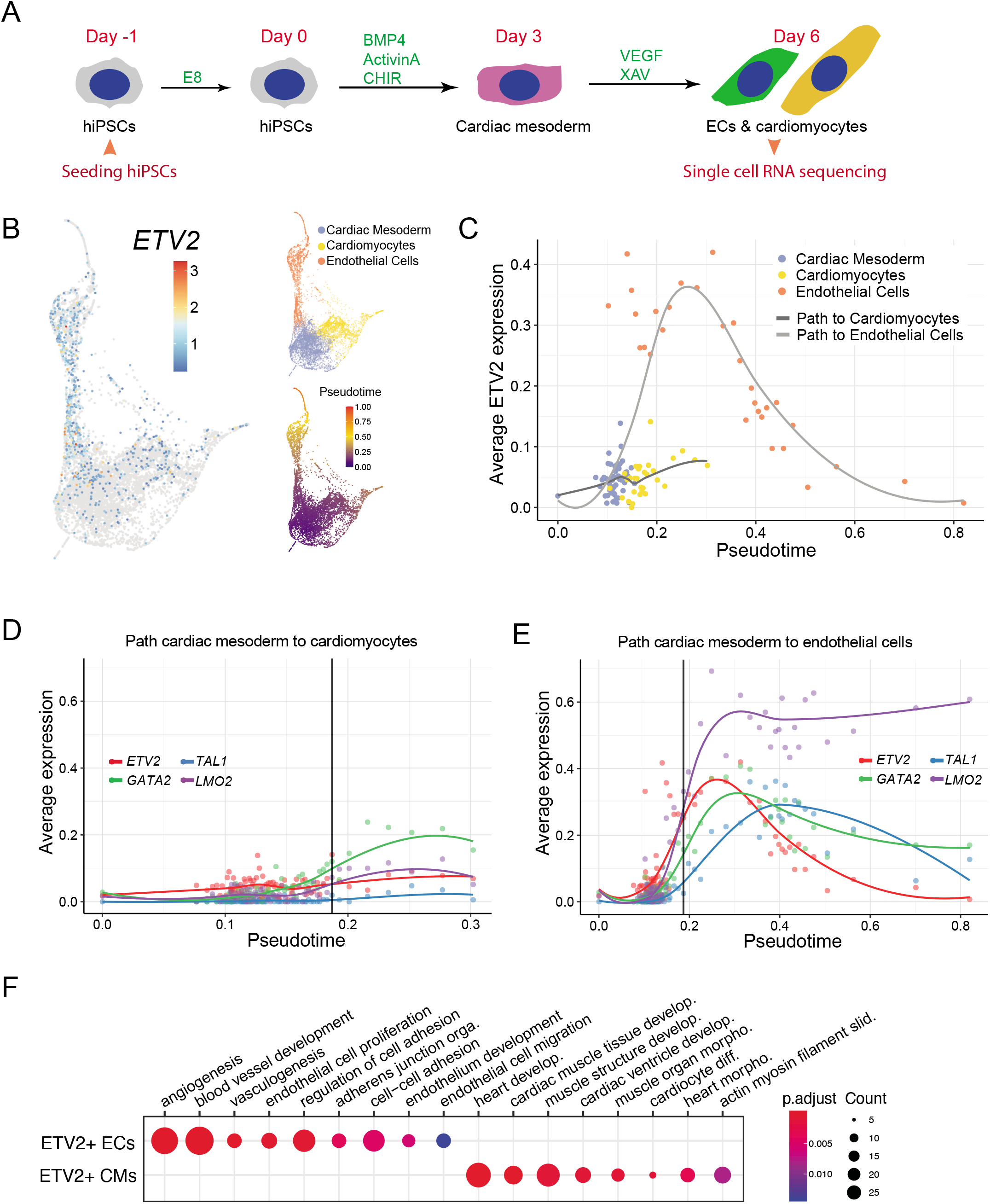
scRNA-seq analysis of ECs and CMs during co-differentiation reveals transient ETV2 upregulation after bifurcation. **(A)** Schematic overview of the co-differentiation protocol from day −1 to day 6. Cells were collected for scRNA-seq on day 6. **(B)** Two-dimensional representation of the scRNA-seq data. Each data point is a single cell. Left: log_2_ transformed *ETV2* expression is indicated by color. Top right: Cell identities are labeled with different colors. Bottom right: Pseudotime is indicated by color. **(C)** Average expression of *ETV2* in bins of pseudotime for the developmental path of CMs or ECs. Cell identities are labeled with different colors. **(D-E)** Average expression of *ETV2* and its direct target genes *TAL1, GATA2, LMO* across binned pseudotime along the developmental path of CMs (D) or ECs (E). Threshold (indicated in black) is set to the timepoint where the average ETV2 expression in EC reaches 0.25. **(F)** GO enrichment analysis of genes that were differentially expressed between ETV2+ ECs and ETV2+ CMs in the scRNA-seq dataset. 128 and 136 genes were upregulated in ETV2+ ECs and CMs respectively (P_adjusted_ < 0.05). A complete list of GO terms can be found in Table S3. Color represents the P_adjusted_ of the enrichment analysis and dot size represents the count of genes mapped to the GO term.

#### Differential expression analysis and identification of cluster maker genes

The R package *edgeR* (V 3.24.3, 31) [42] was used to perform differential expression analysis. We used raw counts and a negative binomial distribution to fit the generalized linear model. The covariates were comprised of 6 binary dummy variables that indicate the three remaining clusters per replicate and a variable that corresponds to the total number of counts per cell. Finally, p-values for each cluster considering both replicates were obtained and adjusted for multiple hypothesis testing with the Benjamini-Hochberg method.

#### Comparison to bulk RNA-sequencing data

The MNN approach was used to integrate the two single-cell replicates, using normalized counts and the 10% HVGs per replicate, and the bulk RNA-sequencing data, with d = 30 and k = 20. After batch correction a diffusion map was calculated on the MNN corrected values with default parameters.

### Statistics

Statistical analysis was conducted with GraphPad Prism 7 software. Data are represented as mean± standard deviation. A Student’s t-test was used for the comparison of two samples. Ordinary one-way ANOVA was used for multiple sample comparison, and uncorrected Fisher’s LSD test was applied. Two-way ANOVA was used for multiple group comparison and uncorrected Fisher’s LSD test was applied. p < 0.05 was considered significant.

## RESULTS

### *ETV2* is upregulated after bifurcation of progenitors into CMs and ECs

To characterize the expression of *ETV2* during co-differentiation of ECs and CMs [33] (Figure 1A), we collected scRNA-seq data on day 6 of differentiation (Figure 1B). We identified three distinct clusters: cardiac mesoderm, CMs and ECs (Figure 1B, top right and Table S3). Pseudotime analysis revealed cardiac mesoderm as the common developmental origin of CMs and ECs (Figure 1B, bottom right). We found that *ETV2* was highly expressed in the EC cluster, as well as in a small fraction of cells in the cardiac mesoderm and CM clusters (Figure 1B, left). We next focused on *ETV2* expression dynamics along the developmental path from cardiac mesoderm to CMs and ECs. Notably, ECs extended to larger pseudotimes (0.15-0.8) compared to CMs (0.15-0.3), which might indicate faster differentiation kinetics in the EC lineage (Figure 1C, S1A). After the bifurcation into ECs and CMs (around pseudotime 0.15),*ETV2* increased only slightly in the CM lineage. In the EC lineage, however, it was initially strongly upregulated (until pseudotime 0.25), and subsequently declined to a similar level as in cardiac mesoderm (Figure 1C). *ETV2* downstream target genes, such as *TAL1, GATA2* and *LMO2* [18], were only slightly increased in the CM lineage (Figure 1D), while in the EC lineage, they were highly induced and strongly correlated with *ETV2* (Figure 1E). Notably, *TAL1, GATA2* and *LMO2* only showed significant expression after ETV2 expression exceeded 0.25 in ECs, an expression level that was not reached in CMs (Figure S1B-C). Endothelial specific genes *KDR, CD34, SOX17, CDH5* and *PECAM1* increased on the path from cardiac mesoderm to ECs (Figure S1E). Most of these genes started to increase when ETV2 was already declining, as exemplified by *CDH5* (Figure S1D). Genes related to cardiac or muscle function, like *ACTC1, PDLIM5, HAND1, PKP2* and *GATA4*, most of which were already expressed in the cardiac mesoderm, were further increased in the CM lineage (Figure S1F). Identification of genes that are differentially expressed between *ETV2+* CMs and ECs showed enrichment in CM- and EC-specific genes, respectively (Figure 1F, Table S4). Taken together, these analyses confirmed the differentiation of cardiac mesoderm into CMs and ECs, which we had discovered previously. They also revealed the increase of *ETV2* as a global indicator of early lineage separation and a transient pulse of high *ETV2* at the beginning of EC specification.

### Generation and characterization of an ETV2^mCherry^ fluorescent hiPSC reporter line

In order to follow *ETV2* expression in real-time, we introduced a fluorescent reporter for *ETV2* in hiPSCs. P2A-mCherry with a nuclear localization signal (NLS) and a neomycin selection cassette was inserted into the endogenous *ETV2* locus before the stop codon using CRISPR/Cas9-facilitated homologous recombination (Figure 2A, S2A). After neomycin selection and excision of the selection cassette, targeted hiPSC clones were validated by PCR (Figure S2A-D) and Sanger sequencing (data not shown). The hiPSC clone with ETV2^mCherry^ in both alleles was further characterized by measuring pluripotency marker expression and G-banding karyotyping (Figure S2E-H). Karyotype analysis revealed an additional duplication in the 1q32.1 locus (Figure S2H). This duplication occurs frequently in hPSCs possibly imposing positive natural selection [53]. However, this did not appear to affect the differentiation of the hPSCs to ECs.

**Figure 2.**
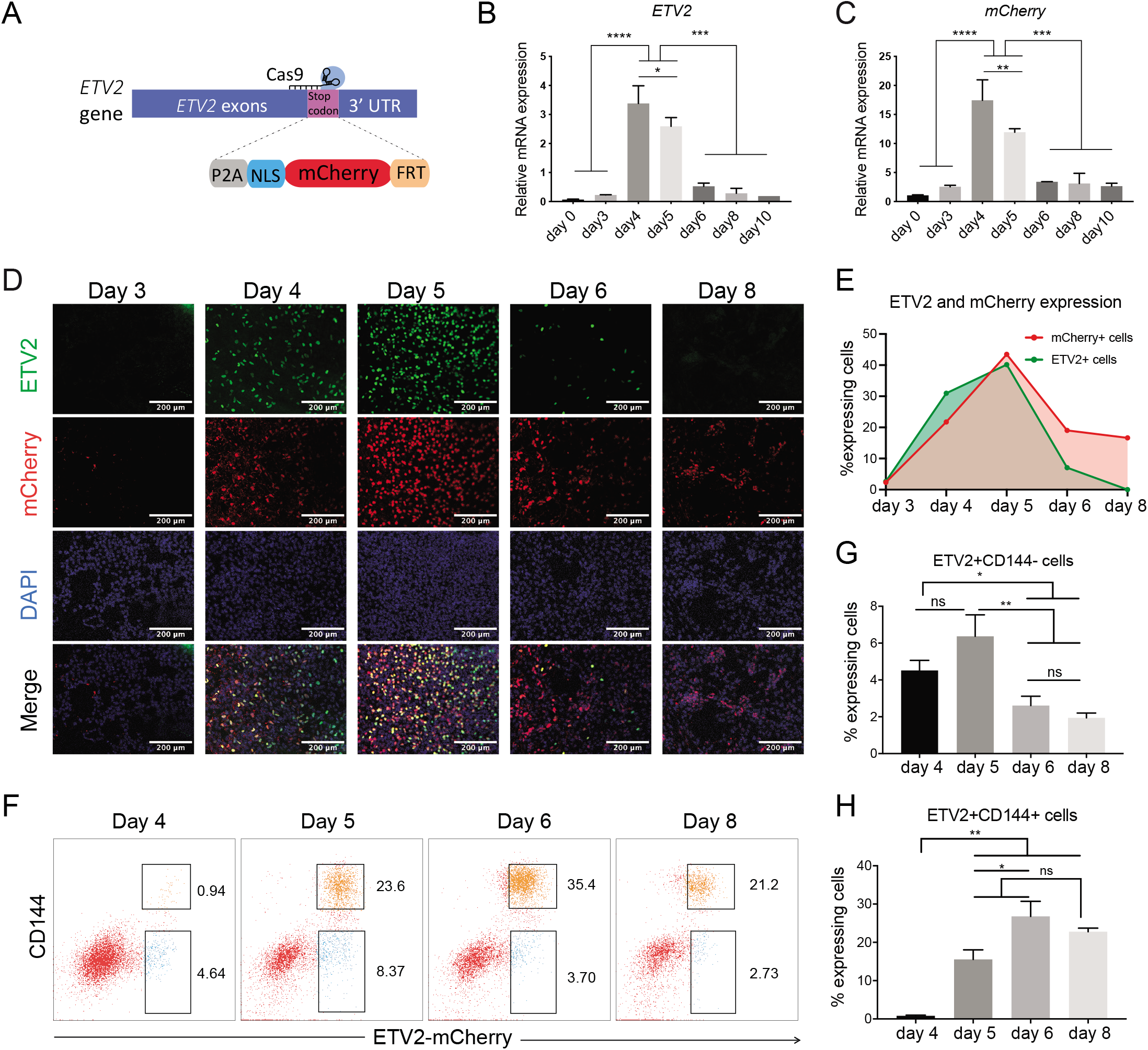
Generation and characterization of an ETV2^mCherry^ hiPSC reporter line. **(A)** Schematic of CRISPR/Cas9-Mediated Knock-in of mCherry into the *ETV2* locus of hiPSCs. **(B-C)** Quantification of *ETV2* and *mCherry* expression by qPCR during differentiation. **(D)** Representative confocal images of ETV2, DAPI and mCherry expression on different days of differentiation. Scale bar 200 μm. **(E)** Quantification of percentage (%) of ETV2+ and mCherry+ nuclei in all DAPI+ nuclei in the field of view in **(D). (F)** Fluorescence activated cell sorting (FACS) based on CD144 and ETV2^mCherry^ expression on day 4, 5, 6 and 8 of differentiation. **(G-H)** Quantification of ETV2+CD144- (SP) ETV2+CD144+ (DP) cells by flow cytometry on differentiation day 4, 5, 6 and 8. (B-C, G-H) Error bars are standard deviations calculated from three independent experiments. Uncorrected Fisher’s LSD test. ns = non-significant, *p < 0.05, **p < 0.01, ***p < 0.001.

*ETV2* and *mCherry* mRNAs were highly expressed on days 4 to 5 of differentiation and downregulated from day 6 (Figure 2B-C). ETV2 and mCherry protein appeared from day 4 and peaked on day 5. ETV2 protein was downregulated on day 6 and absent on day 8. mCherry was retained in a fraction of the cell population for somewhat longer because of its half-life (Figure 2D-E, S3A and supplemental online Video 1). Flow cytometry analysis at different stages of differentiation revealed upregulation of ETV2 (mCherry protein) as early as day 4 of differentiation followed by the upregulation of the EC-specific marker CD144 (Figure 2F, S3B). Quantification of the percentages of single positive (SP; ETV2^mCherry^+CD144-) and double positive (DP; ETV2^mCherry^+CD144+) cells on day 4, 5, 6 and 8 of differentiation from at least three independent experiments showed a decrease and an increase of SP and DP cells respectively (Figure 2G-H). mCherry protein remained present for a longer period than ETV2 protein and endogenous *ETV2 and mCherry* mRNA (Figure 2B-H), likely due to the relatively longer half-life of the mCherry protein. This explains the persistence of mCherry signal in both the DP and SP population (Figure 2G-H), and offers the possibility to use mCherry as a lineage tracer, identifying cells that previously passed through a stage of being ETV2+.

### The ETV2^mCherry^ fluorescent reporter allows the purification of differentiating cells with lineage-specific expression profiles

We next sorted DP and SP cells at different stages of differentiation (Figure 2F) and performed bulk RNA-seq on at least three independent replicates. *ETV2* mRNA showed similar trends in DP and SP cells (Figure S4), consistent with the earlier qPCR result (Figure 2B).

Principal component analysis (PCA) showed that DP and SP populations diverged progressively over time (Figure 3A). Mapping of the bulk transcriptomes to the scRNA-seq data revealed that DP samples aligned on the EC branch and SP cells on the CM branch (Figure 3B). Notably, SP and DP cells collected at later time points were further away from cardiac mesoderm, reflecting ongoing differentiation (Figure 3B).

**Figure 3.**
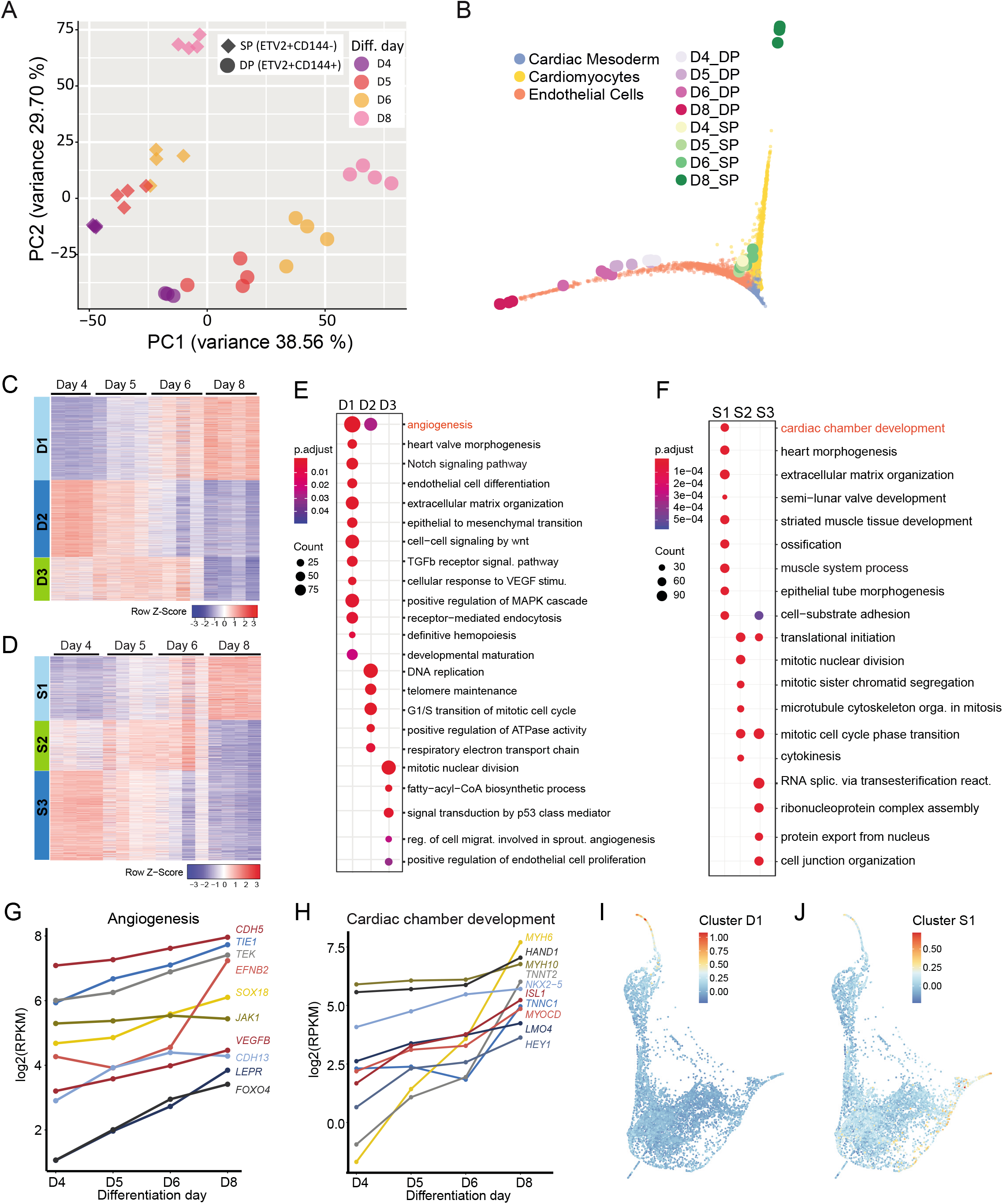
Bulk RNA-seq of the ETV2^mCherry^ reporter line shows diverging transcriptional profiles. **(A)** PCA of all sorted DP and SP samples collected from three or four independent differentiations. **(B)** Low-dimensional representation (diffusion map) of scRNA-seq and bulk RNA-seq samples collected on day 4, 5, 6 and 8. The small data points correspond to individual cells, the large symbols correspond to bulk samples. Different clusters of cells or bulk samples are labelled with different colors. **(C-D)** Gene expression pattern in all DP (D) and SP (E) cells. The 3000 most variable genes across all DP or SP samples were identified and grouped into three clusters by consensus clustering. The genes in each cluster can be found in Table S5. The color scale represents relative expression (row-wise z-score). **(E-F)** GO enrichment analysis of each gene cluster of DP (E) and SP (F) samples. Representative GO terms are shown. The complete list of GO terms can be found in Table S6. Color represents the P_adjusted_ of the enrichment analysis and dot size represents the count of genes mapped to the GO term. **(G-H)** Representative genes mapped to representative GO terms of clusters D1 (G) and S1 (H) and their expression levels from day 4 to day 8 are shown. **(I-J)** Low-dimensional representation of the scRNA-seq data. Each data point is a single cell. Mean expression of genes in cluster D1 (I) and S1 (J) in the scRNA-seq data is indicated by color. Gene expression was scaled gene-wise prior to averaging.

We next leveraged the higher sensitivity and accuracy of bulk RNA-seq compared to scRNA-seq, to get a more comprehensive and robust transcriptional characterization of the subpopulations. By consensus clustering of the most variable genes across DP or SP cells (3000 genes each) we found three gene clusters for each population, with distinct expression dynamics (Figure 3C-D, Figure S5A-B, Table S5). In DP cells, cluster D1 (1226 genes) expression increased over time. Gene Ontology (GO) terms enriched in cluster D1 included “angiogenesis”, “Notch signaling pathway”, “transforming growth factor beta receptor signaling pathway”, “receptor-mediated endocytosis” and “developmental maturation” (Figure 3E, Table S6). In accordance with this analysis, angiogenesis-related genes (*CDH5, TIE1, TEK, EFNB2, SOX18, VEGFB, LEPR*), Notch and transforming growth factor beta receptor signaling pathway related genes (*COL1A2, NOTCH1, HES4, DLL4, JAG2, HEY1, NOTCH3, NOTCH4, TGFBR2, EGF*) and heart valve morphogenesis related genes (*SMAD6, EFNA1, GATA5, HEY2, EMILIN1, NOS3, GATA3*) were upregulated over the course of differentiation in DP cells (Figure 3G, Figure S5C). In the scRNA-seq data, cluster D1 genes were specifically expressed in the EC cluster and showed increasing expression along pseudotime (Figure 3I). Cluster D1 genes are thus likely involved in EC-specific functions. Cluster D2 (1127 genes), which was downregulated after day 4 (Figure 3C), was enriched for cell cycle-related genes (*ITGB1, CDK4, CCND1, CDK2AP2, MYC, CDC6*)(Figure S5D). Cluster D3 (647 genes), which was downregulated after day 5-6 (Figure 3C), contained cell proliferation- and fatty-acyl-CoA biosynthetic process-related genes (*ACLY*, *FASN, ELOVL1, SLC25A1, ACSL3, ACSL4*)(Figure S5E). Genes in clusters D2 and D3 were more broadly expressed in the scRNA-seq data (Figure S5I). Their dynamics likely reflect changes in proliferation and metabolism at the exit from the multipotent progenitor state.

In SP cells, cluster S1 (936 genes) increased over time and contained genes enriched for GO terms related to heart development and function (Figure 3F, Table S6). In agreement, cardiac chamber and cardiac muscle development related genes (*MYH6, HAND1, MYH10, TNNT2, NKX2-5, ISL1, TNNC1, MYOD, LMO4* and *HEY1, MYL7, MYL4, ACTA2, KCNH2*) were upregulated over the course of differentiation (Figure 3H, S5F). Cluster S1 genes were highly expressed in the cardiac mesoderm and CM clusters in the scRNA-seq data, which showed an increase over pseudotime in the CM lineage (Figure 3J). These genes are thus likely involved in CM-specific functions. Cluster S2 (746 genes), which increased slightly until day 6 and was downregulated afterwards (Figure 3D), contained mitotic nuclear division genes (TPX2, CDC20, NEK2, PLK1, PRC1 and CDC25C) (Figure S5G). Cluster S3 (1318 genes), whose expression decreased continuously over time (Figure 3D), contained transcription and translation process-related genes (*SF1, SNRPE, DDX23, RRP1B* and *PRMT5*) (Figure S5H). In the scRNA-seq data, genes from clusters S2 and S3 showed broader expression patterns compared to cluster S1 genes (Figure S5J). The dynamics of clusters S2 and S3 likely reflect changes in proliferation and metabolism in the CM lineage, analogous to the role of clusters D2 and D3 in the EC lineage.

Taken together, time-resolved bulk RNA-seq of sorted SP and DP populations confirmed that ETV2-positive cells contained transcriptionally distinct subpopulations. DP cells were part of the EC lineage, while SP cells corresponded to the CM lineage.

### ETV2+ cells contain lineage-predisposed subpopulations

Next, we wanted to find out how the various subpopulations we identified differed in terms of their further differentiation potential. To this end, we sorted cells on the basis of ETV2 reporter levels shortly after the bifurcation (on day 5) and attempted to differentiate them further towards ECs by adding VEGF (Figure 4A-B). After 5 days of additional differentiation, ETV2+ cells produced more than 90% CD144+CD31+ ECs, while ETV2-cells gave rise to only 10-15% ECs (Figure 4C-D, S6A). Only cells derived from ETV2+ cells expressed endothelial-specific markers, as observed by qPCR and immunofluorescence (Figure 4E-H, S6C). These cells also upregulated pro-inflammatory markers, such as ICAM-1 and E-Selectin upon TNF-a stimulation (Figure 4I-L, S6B), as shown previously for hiPSC-derived ECs [54]. We thus concluded that the majority of ETV2+ cells on day 5 has a strong propensity to produce ECs.

**Figure 4.**
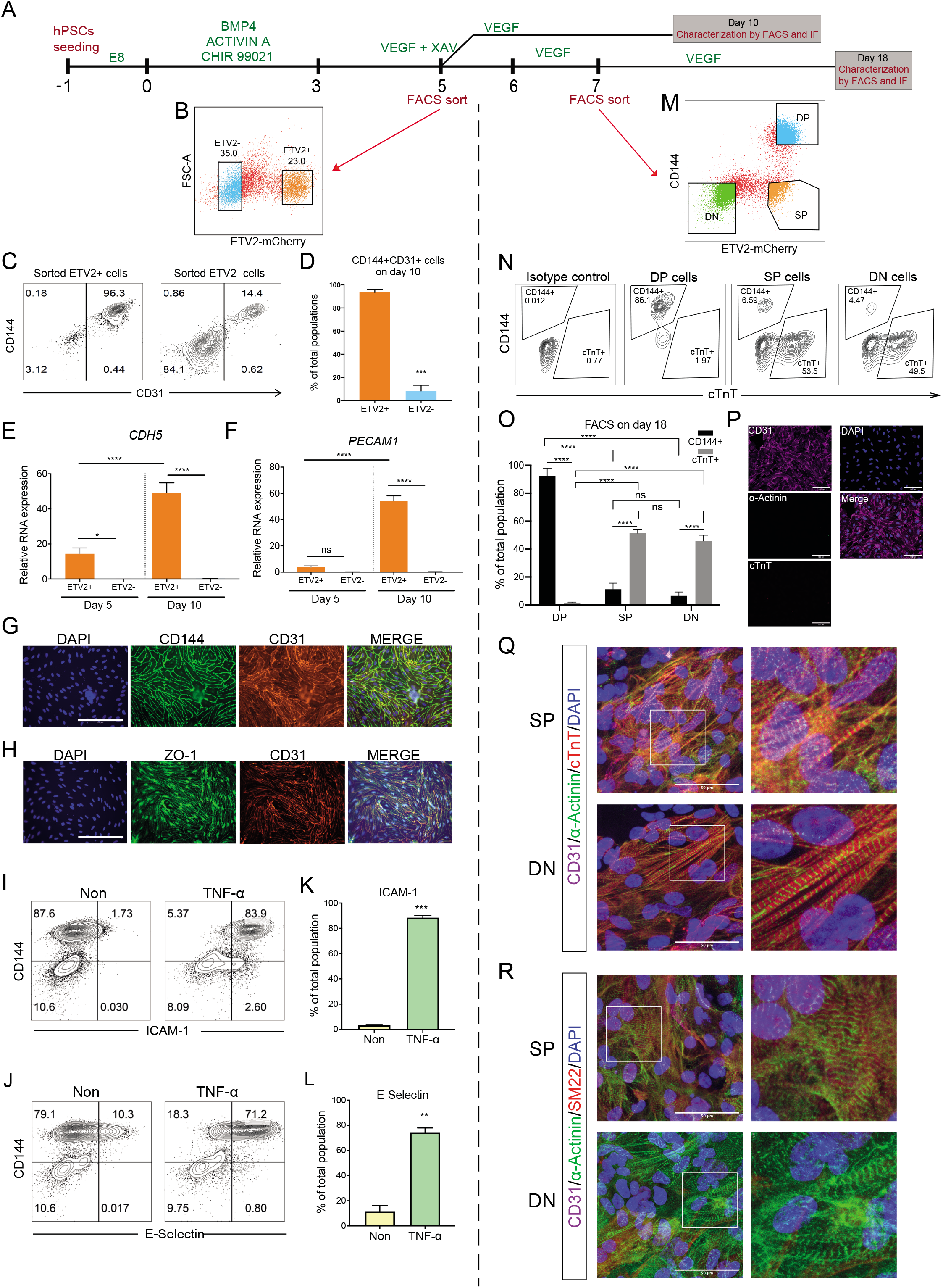
ETV2+ cells contain two lineage-predisposed subpopulations. **(A)** Schematic of the differentiation protocol and cell sorting. ETV+ and ETV2-cells were sorted on day 5 and cultured in VEGF until day 10. DP, SP, DN were sorted on day 7 and cultured in VEGF until day 18. **(B)** Representative flow cytometry analysis ETV2-mCherry expression on day 5 and gates for cell sorting of ETV2+ and ETV2-cells are shown. **(C)** Flow cytometry analysis of endothelial markers CD144 and CD31 on day 10 of sorted ETV2+ and ETV2-cell differentiation. **(D)** Quantification of CD144+CD31+ cells in the total population on day 10 of sorted ETV2+ and ETV2-cell differentiation. **(E-F)** Quantification of *CDH5* and *PECAM1* expression in sorted ETV2+ and ETV2-cells on day 5 and day 10. **(G-H)** Immuno-staining of CD144, CD31 and cell-cell junctional marker ZO-1 on day 10 for sorted ETV2+ cells. Scale bar 200 μm. **(I-J)** Flow cytometry analysis of ICAM1, E-Selectin and CD144 for sorted ETV2+ cells on day 10. Cells were stimulated with TNF-α for 24 h before analysis. **(K-L)** Quantification of CD144+ICAM-1+ (K) and CD144+E-Selectin+ (L) cells in the population on day 10. **(M)** Flow cytometry analysis of CD144 and ETV2-mCherry expression on day 7. DP, SP and DN cells were gated and sorted. **(N)** Flow cytometry analysis of CD144 and CM marker cTnT expression on day 18 of sorted DP, SP and DN cells. Isotype control antibodies were included as negative control. **(O)** Quantification of CD144+ ECs and cTnT+ CMs on day 18 of DP, SP and DN cell differentiation. **(P)** Immuno-staining of CD31, a-Actinin, cTnT and DAPI on day 18 of DP cell differentiation. Scale bar 50 μm. **(Q-R)** Immuno-staining of CD31, a-Actinin, cTnT, SM22 and DAPI on day 18 of SP and DN cells. Scale bar 50 μm. Error bars are ± SD of three independent experiments in (D-F, K-L, O). T test (D, K, L) and uncorrected Fisher’s LSD test (E-F, O) were used. ns = non-significant, *p < 0.05, **p < 0.01, ***p < 0.001, ****p < 0.0001.

Both the analysis of the scRNA-seq data and the time-resolved bulk RNA-seq of sorted cells had identified a subpopulation of ETV2+ cells with CM characteristics. We strongly suspected that the differentiation of these cells would be predisposed to the CM lineage. To test this hypothesis with our reporter line, we co-differentiated cells until day 7. We chose a later time point for this experiment because the majority of cells are past bifurcation at this point and it is therefore easier to identify the ETV2+ population that does not correspond to early ECs. We co-stained for CD144 and sorted the cells into DP, SP and double negative (DN) populations. These subpopulations were then further cultured in the presence of VEGF until day 18 (Figure 4A, M). The majority (>80%) of DP cells differentiated into CD144+CD31+ ECs, in agreement with the previous experiment (Figure 4N-O). In contrast, more than 50% of SP and DN cells differentiated into cTnT+ CMs while very few ECs were detected (Figure 4N-O). Interestingly, CMs derived from SP cells seemed to proliferate more and formed a monolayer composed of a contracting cell sheet, while CMs from DN cells proliferated to a lesser extent and produced only a few, isolated clusters of contracting cells (supplemental online Video 2). Almost all DP cells on day 18 expressed the EC marker CD31, while only few cells derived from SP and DN cells were positive for CD31 (Figure 4P-R). Most cells derived from SP and DN expressed CM-specific α-Actinin and cTnT and showed typical sarcomeric structures (Figure 4Q-R, S6D-E). A small number of SP and DN-derived cells were also positive for the smooth muscle cell marker SM22, while negative for cardiac markers (data not shown). Furthermore, the α-Actinin positive CMs derived from the SP cell fraction were positive for SM22, possibly indicating their immaturity (Figure 4R).

Taken together, the VEGF differentiation experiments showed that DP and SP cells are predisposed to the EC and CM lineages, respectively. DN cells were largely unable to give rise to EC but produced CMs, albeit with lower efficiency than SP cells. Entering a transient state characterized by high ETV2 expression, thus seems necessary to initiate EC specification.

## DISCUSSION

In this study, we characterized the dynamics of EC and CM co-differentiation from hiPSCs [33]. *ETV2* was identified as an early indicator of lineage segregation and found to be strongly, but transiently, upregulated in ECs, in agreement with its essential role in hemangiogenic development [55]. Interestingly, *ETV2* expression was also observed in a small population of cardiac mesoderm and CMs. This is reminiscent of a recent study where etv2 expression was observed in lateral plate mesoderm and the CM population in zebrafish [56]. In our experiments, expression of ETV2 target genes seemed to occur only above a threshold of *ETV2* expression, although this observation could also be explained by a temporal delay between ETV2 upregulation and target gene expression. An ETV2 threshold in hiPSC differentiation would be in line with previous reports of an ETV2 threshold in hemangiogenic specification [19,20]. Our results thus support an *ETV2* pulse- and threshold dependent specification of ECs.

With the ETV2^mCherry^ hiPSC reporter line, generated to track, isolate and characterize *ETV2+* cells, we showed that ETV2+ cells could give rise to both ECs and CMs. Over time, EC and CM precursors acquired more specific endothelial and myocardial identities, respectively, as well as downregulating cell cycle-related genes, which indicated exit from the progenitor state and further maturation.

In the DP subpopulation (EC precursors), several key angiogenesis and Notch signaling pathway genes, like *LEPR, FOXO4, DLL4, NOTCH4* and *EGF*, strongly increased starting from day 4, indicating a specified EC fate but an immature state on day 4. These relatively late expressed genes could potentially be used as markers to distinguish early and late ECs during development *in vitro* or *in vivo*.Genes involved in heart development and definitive hematopoiesis were also upregulated during EC development, suggesting a mixture of cardiac endothelial- and hemogenic endothelial identity of these ECs. A better characterization hematopoietic potential of these cells would be interesting but beyond the scope of this study. ECs that were further differentiated with VEGF showed a clear endothelial identity and were fully functional based on their inflammatory response upon TNFa stimulation. Notably, they also expressed a number of cardiac markers like *MEOX2, GATA4, GATA6* and *ISL1*, suggesting a cardiac specific EC identity [33].

The SP subpopulation (CM precursors) had already committed to a cardiac fate on day 4, as evidenced by expression of cardiac genes *HAND1, MYH10, NKX2-5, ISL1, TNNC1, MYOCD* and *LMO4*.However, some crucial CM genes were still absent, including *MYH6* and *TNNT2. MYH6* encodes the major CM thick filament protein MHC-α and *TNNT2* is routinely used as a CM marker. Both genes are essential for CM contractility and started to be expressed only after day 4. Their relatively late expression could allow us to identify early and late cardiac progenitors during cardiac development in future studies. CMs were still early progenitors on day 6 of the differentiation as no functionally contracting CMs were observed yet at this stage. Pseudotime analysis also suggested that ECs had differentiated further compared to CMs on day 6. After additional VEGF differentiation, SP cells gave rise to contracting CM, which provided direct evidence they were CM precursors. More importantly, it demonstrated that both ECs and CMs could be derived from ETV2+ progenitors, confirming the presence of a common precursor implied by our earlier studies [33].

Notably, ETV2-cells (DN population) also gave rise to contracting CMs after VEGF treatment, albeit less frequently than SP cells. This difference could be due to either the different cell growth rates or their different developmental origins (FHF vs. SHF). More work is needed to establish the identity of CMs from SP and DN populations in the future.

## CONCLUSION

Bulk- and single cell transcriptomic analysis in this study provide insight into the differentiation dynamics of cardiomyocytes and cardiac endothelial cells, two important human cardiac lineages. This rich data set is now available for comparison with *in vivo* data. The ETV2 fluorescent reporter we generated in hiPSCs allowed identification of a new subpopulation of early CM precursors that expressed ETV2.

## ACKNOWLEDGEMENTS

S.L. Kloet and E. de Meijer (Leiden Genome Technology Center) for help with 10X Genomics experiments (cell encapsulation, library preparation, single-cell sequencing, primary data mapping, and quality control). K. Anastassiadis (Technical University Dresden) for providing P15 backbone with a Neomycin resistance cassette surrounded by two FRT sequences and CAGGs-Flpo-IRES-puro vector. R. Davis (Leiden University Medical Center) for comments on the manuscript. B. van Meer for input into the realtime imaging using the Incucyte^®^ system. Sartorius Stedim Biotech GmbH for usage of the Incucyte^®^ Live-Cell Analysis System.

## Data Availability Statement

The accession numbers for the bulk and single cell RNA sequencing datasets reported in this paper are https://www.ncbi.nlm.nih.gov/geo/ GEO: GSE157954 (bulk) and GEO: GSE202901 (single cell).

---

**Figure S1.**
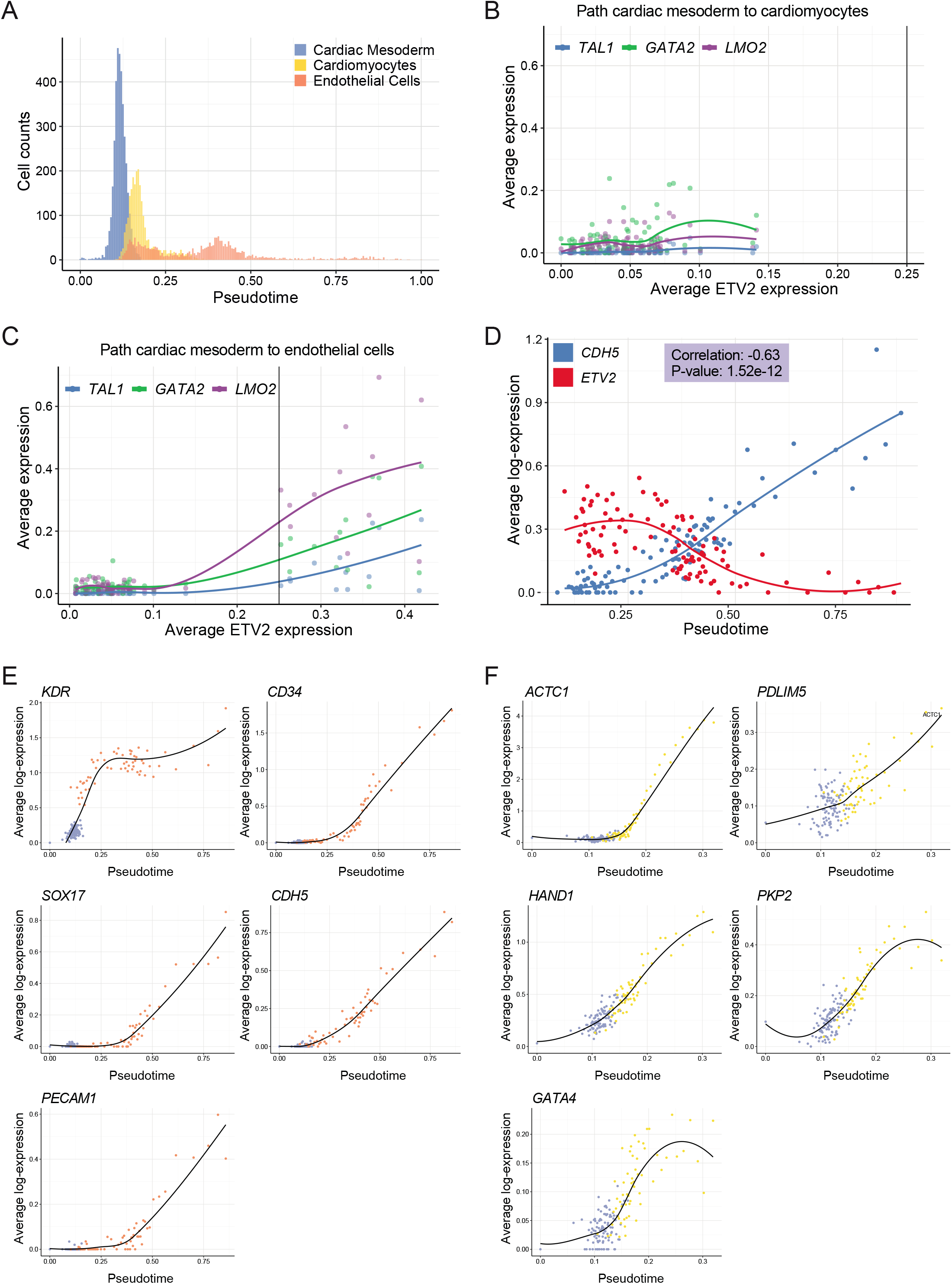
Pseudotime analysis of EC and CM co-differentiation. **(A)** Distribution in pseudotime for each cell cluster. **(B-C)** Average expression of *ETV2* and *ETV2* target genes along the developmental path of CMs (B) and ECs (C). The vertical black line indicates an *ETV2* expression level of 0.25. **(D)** Average expression of *CDH5* and *ETV2* in the EC cluster across pseudotime. Binning and averaging were performed as for (B) and (C). The p-value for the correlation between *CDH5* and *ETV2* expression is based on the null hypothesis that the correlation is zero. **(E)** Expression of endothelial markers across pseudotime during the development path of ECs. **(F)** Expression of cardiac markers across pseudotime during the development path of CMs.

**Figure S2.**
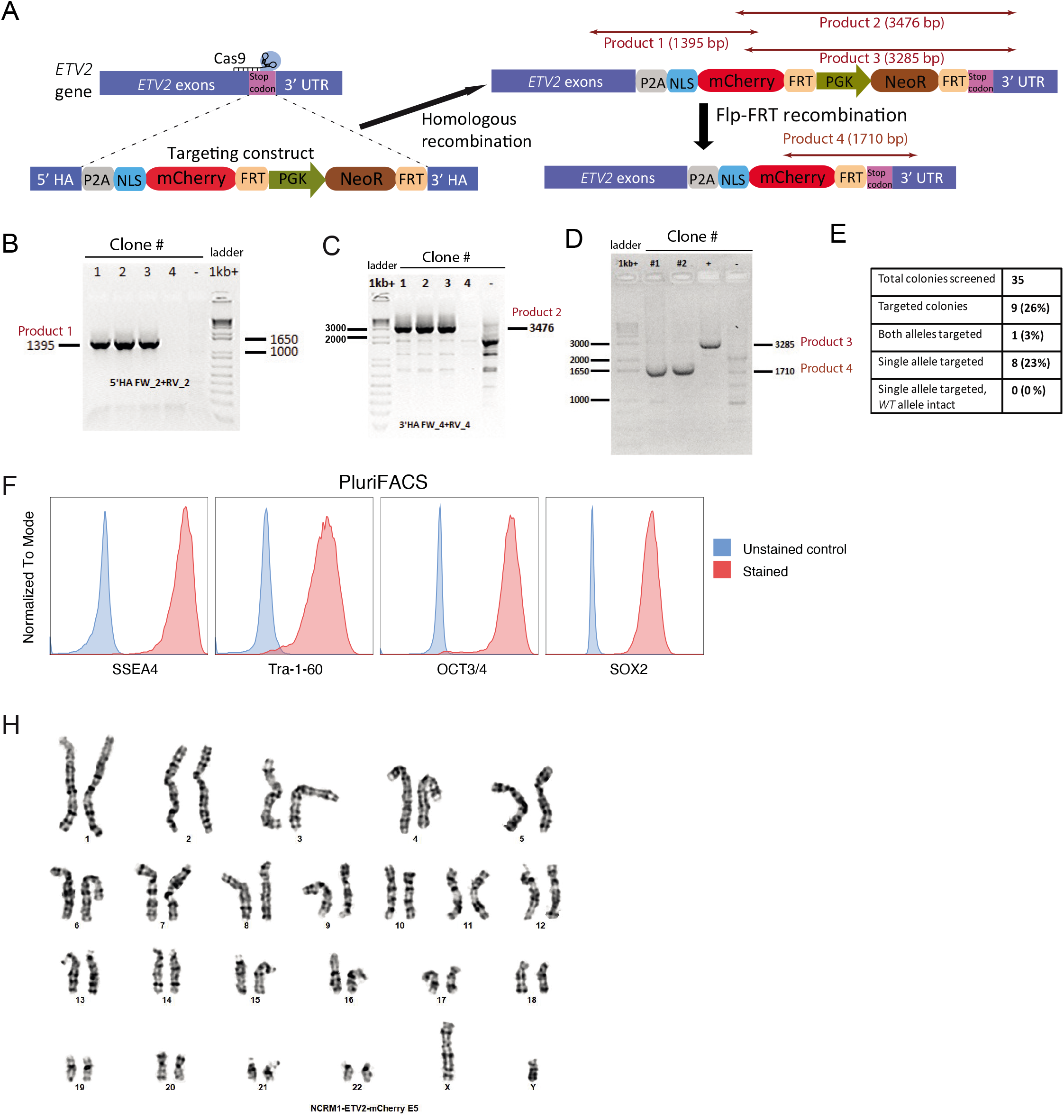
Generation and characterization of ETV2^mCherry^ hiPSC reporter line. **(A)** Schematic of CRISPR/Cas9-Mediated Knock-in of mCherry into the *ETV2* locus of hiPSCs. mCherry and Neomycin Resistance (NeoR) sequences were inserted into the *ETV2* locus through homologous recombination. Then NeoR cassette was removed by flpo recombinase. 4 pairs of primers were used for PCR screening of targeted clones. **(B-C)** PCR screening of targeted clones with correct insertion at the *ETV2* locus. Two pairs of primers (for product 1 and 2) were used to confirm the integration of the construct. Clone 1, 2, 3 were correctly targeted and clone 4 was not targeted. Non-targeted hiPSCs (-) were included as negative control. **(D)** The excision of the neomycin-resistance cassette was confirmed by PCR (product 3 and 4 are present before and after excision, respectively). Clones 1 and 2 were successfully excised. Genomic DNA before excision (+) and non-targeted hiPSCs (-) were included as positive and negative control, respectively. **(E)** Summary of CRISPR targeting efficiency. Of 35 colonies screened, 1 colony was targeted in both alleles. 8 colonies were targeted in only one allele but the other allele showed unwanted mutations. **(F)** Flow cytometry analysis of pluripotency markers in targeted ETV2^mCherry^ hiPSC clone. **(H)** A representative karyogram (G-banding) of targeted ETV2^mCherry^ hiPSC clone.

**Figure S3.**
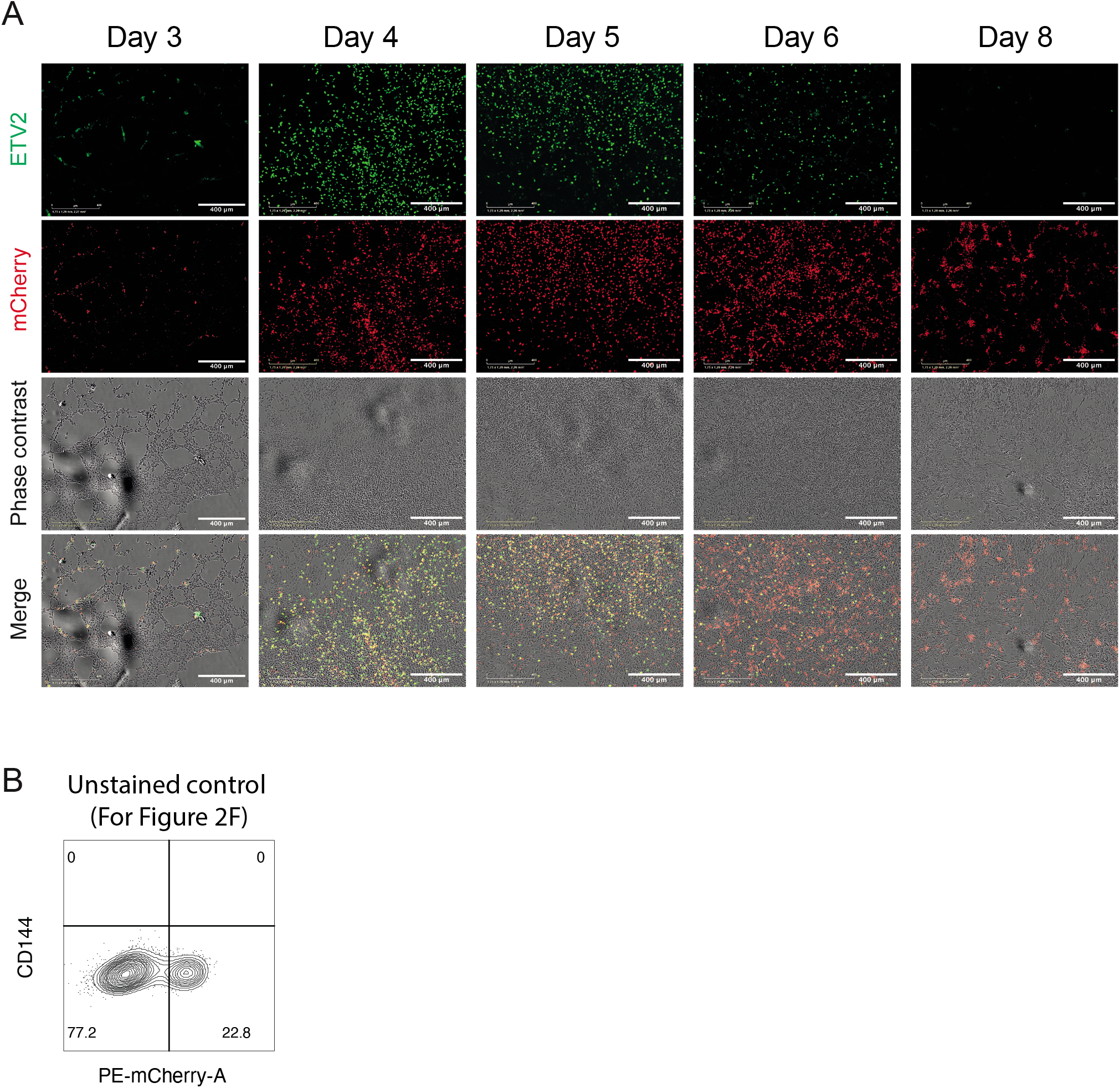
Characterization of an ETV2^mCherry^ hiPSC reporter line. **(A)** Immuno-staining of ETV2, mCherry expression and phase contrast were imaged on the Incucyte system on different days of differentiation. Scale bar 400 μm. **(B)** Unstained control for the FACS analysis shown in Figure 2F.

**Figure S4.**
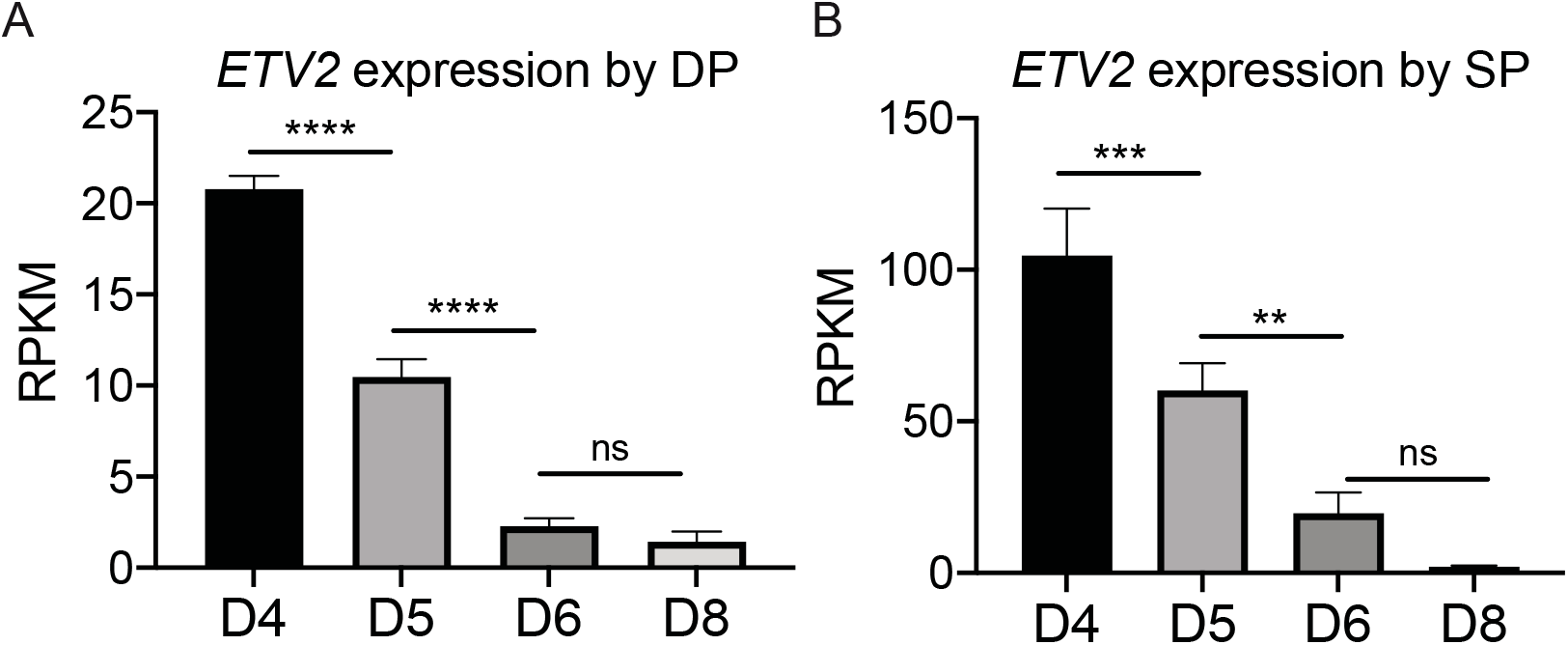
*ETV2* expression in DP and SP cells during EC and CM co-differentiation. **(A-B)** Normalized expression value (RPKM) of *ETV2* in DP **(A)** and SP **(B)** cells sorted on day 4, 5, 6 and 8. Error bars are ± SD of four independent experiments. Uncorrected Fisher’s LSD test. ns = non-significant, *p < 0.05, **p < 0.01, ***p < 0.001, ****p < 0.0001.

**Figure S5.**
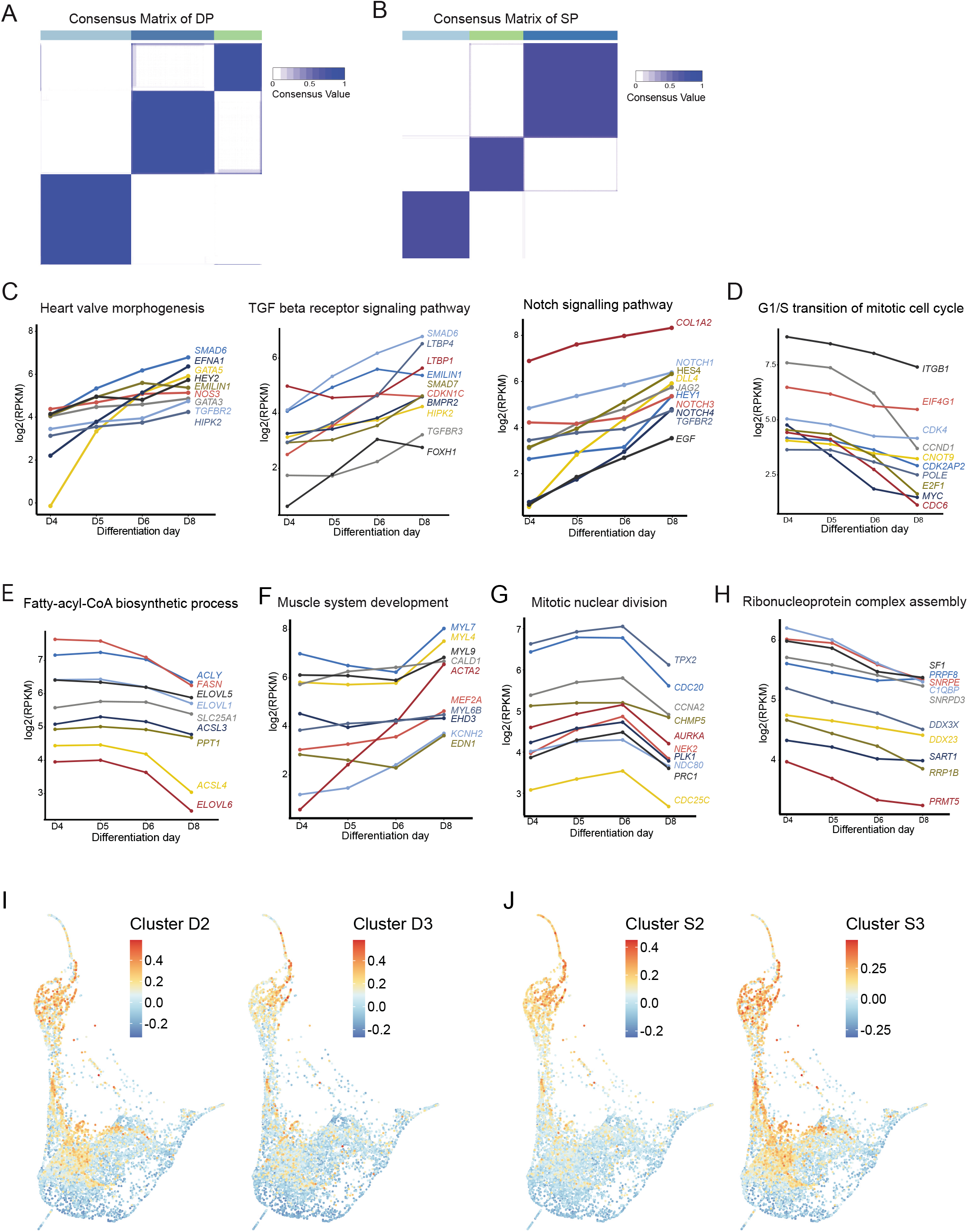
Bulk RNA-seq analysis of EC and CM differentiation. **(A-B)** Consensus clustering of the 3000 most variant genes across all DP samples (A) and all SP samples (B). All genes were divided into 3 clusters D1-D3 for DP (A) and S1-S3 for SP (B). Consensus value indicates similarity between two genes. **(C-H)** Representative GO terms enriched in cluster D1 (C), D2 (D), D3 (E), S1 (F), S2 (G) and S3 (H). Representative genes mapped to these GO terms and their expression levels from day 4 to day 8 are shown. **(I-J)** Low-dimensional representation of the scRNA-seq data. Each data point is a single cell. Mean expression of genes in cluster D1 (I) and S1 (J) in the scRNA-seq data is indicated by color. Gene expression was scaled gene-wise prior to averaging.

**Figure S6.**
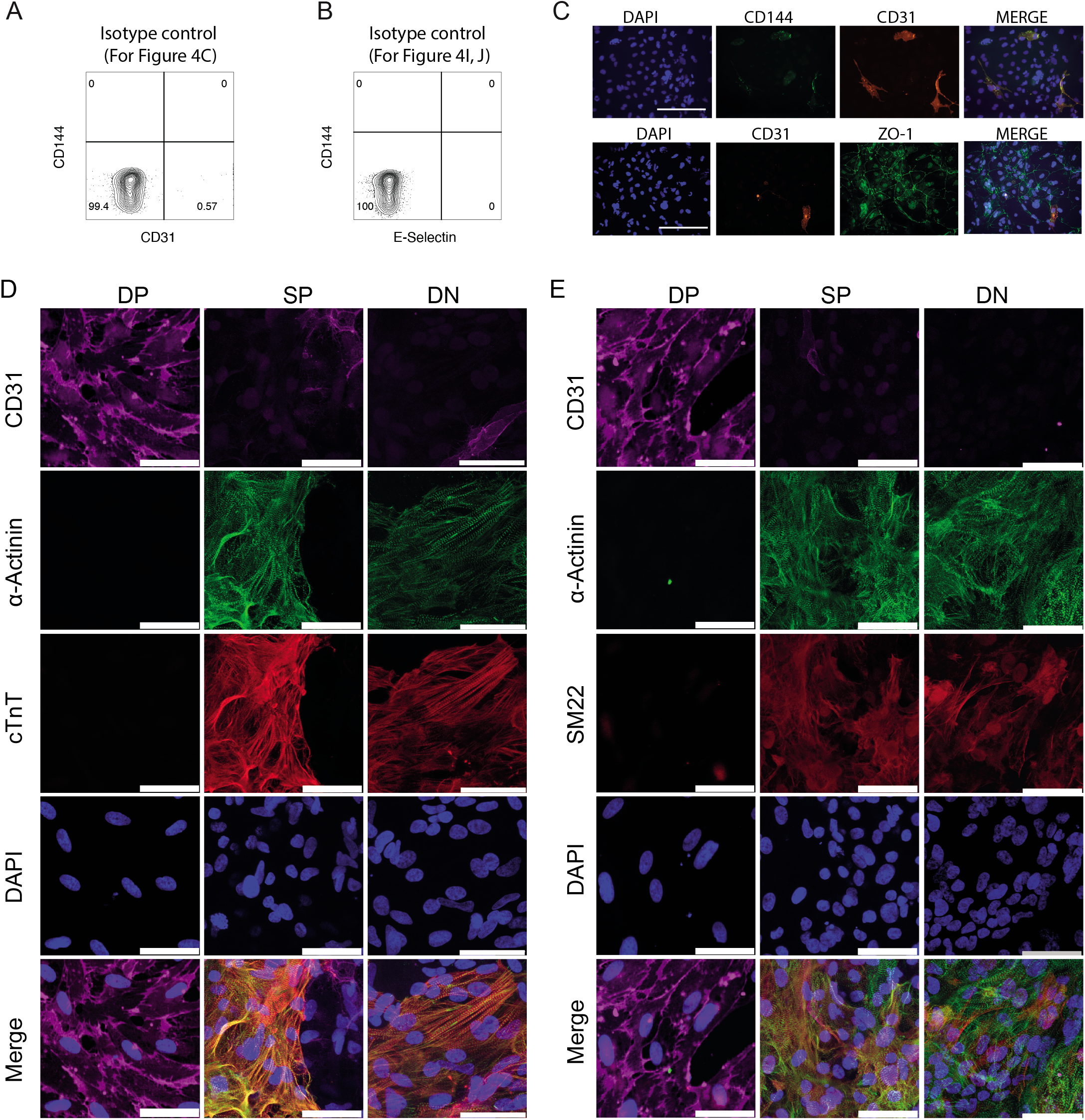
Experimental control for flow cytometry analysis and immuno-staining. **(A-B)** Isotype control for the FACS analysis shown in Figure 4C **(A)** and Figure I-J **(B)**. **(C)** Immuno-staining of CD144, CD31 and cell-cell junctional marker ZO-1 on day 10 for sorted ETV2-cells. Scale bar 200 μm. **(E-F)** Immuno-staining of CD31, a-Actinin, cTnT, SM22 and DAPI on day 18 of DP, SP and DN cells. Scale bar 50 μm.

## SUPPLEMENTARY INFORMATION

### Online supplementary files

Supplemental figures S1-S6 (uploaded as individual files)

Supplemental tables S1-S6 (uploaded as individual files):

Table S1. List of antibodies

Table S2. Sequence of primes used for qPCR

Table S3 Markers for each scRNA-seq cluster

Table S4 List of GO terms for ETV2 expressing cells in EC and CM clusters

Table S5 Gene list in all clusters of SP and DP samples

Table S6 List of GO terms for DP and SP clusters

Supplemental online Video 1 and 2 (uploaded as individual files):

Video 1 Time lapse of differentiation from day 3.5 to day 8.5. 20 min per frame

Video 2 SP and DN cells on day 18 of differentiation

