## Supplementary material for "ETV2 upregulation marks the specification of early cardiomyocytes and endothelial cells during co-differentiation": Table S1

| Antibody | Application | Source | Dilution | Catalog # |
| --- | --- | --- | --- | --- |
| SSEA4 | FACS | BD Biosciences | 1:3200 | 560126 |
| OCT3/4 | FACS | BD Biosciences | 1:50 | 560794 |
| SOX2 | FACS | BD Biosciences | 1:100 | 51-9006407 |
| TRA-1-60 | FACS | BD Biosciences | 1:200 | 563188 |
| CD144 | FACS | Invitrogen | 1:50 | 53-1449-42 |
| ICAM1 | FACS | R&D Systems | 1:20 | BBA20 |
| E-Selectin | FACS | R&D Systems | 1:20 | BBA21 |
| SSEA4 | IF | BioLegend | 1:200 | 330402 |
| OCT3/4 | IF | Santa Cruz | 1:100 | SC-5279 |
| Nanog | IF | Cell Signaling | 1:200 | 4903S |
| ETV2 (rabbit) | IF | Abcam | 1:500 | Ab181847 |
| mCherry (rat) | IF | Invitrogen | 1:500 | M11217 |
| α-ACTININ (mouse) | IF | Sigma-Aldrich | 1:800 | A7811 |
| cTnT (rabbit) | IF | Abcam | 1:500 | ab45932 |
| CD31 (sheep) | IF | R&D | 1:200 | AF806 |
| ZO-1 (rabbit) | IF | Invitrogen | 1:200 | 61-7300 |
| CD144 (rabbit) | IF | Cell Signaling | 1:200 | 2158 |
| SM22 (rabbit) | IF | Abcam | 1:400 | ab14106 |
| AF488 (goat anti-mouse) | IF | Invitrogen | 1:250 | A21151 |
| AF555 (donkey anit-rabbit) | IF | Invitrogen | 1:250 | A31572 |
| AF647 (goat anti-mouse) | IF | Invitrogen | 1:250 | A21242 |
| AF488 (donkey anti-mouse) | IF | Invitrogen | 1:300 | A21042 |
| AF568 (donkey anti-sheep) | IF | Invitrogen | 1:300 | A21099 |
| AF647 (donkey anti-rabbit) | IF | Invitrogen | 1:300 | A31573 |
| AF594 (donkey anti-rat) | IF | Invitrogen | 1:300 | A48271 |
| AF488 (donkey anti-rabbit) | IF | Invitrogen | 1:300 | A21206 |

**Supplemental Table S1. List of antibodies**
