## Supplementary material for "ETV2 upregulation marks the specification of early cardiomyocytes and endothelial cells during co-differentiation": Table S2

| Gene | Forward sequence | Reverse sequence | Product size |
| --- | --- | --- | --- |
| *hARP* | CACCATTGAAATCCTGAGTGATGT | TGACCAGCCCAAAGGAGAAG | 116 |
| *RPL37A* | GTGGTTCCTGCATGAAGACAGTG | TTCTGATGGCGGACTTTACCG | 84 |
| *ETV2* | CAGCTCTCACCGTTTGCTC | AGGAACTGCCACAGCTGAAT | 106 |
| *CDH5* | GGCATCATCAAGCCCATGAA | TCATGTATCGGAGGTCGATGGT | 100 |
| *CD31* | GCATCGTGGTCAACATAACAGAA | GATGGAGCAGGACAGGTTCAG | 101 |
| *PDGFRA* | ATTGCGGAATAACATCGGAG | GCTCAGCCCTGTGAGAAGAC | 95 |
| *NKX2-5* | TTCCCGCCGCCCCCGCCTTCTAT | CGCTCCGCGTTGTCCGCCTCTGT | 138 |
| *TBX5* | ACATGGAGCTGCACAGAATG | TGCTGAAAGGACTGTGGTTG | 104 |
| *GATA4* | GACAATCTGGTTAGGGGAAGC | GAGAGATGCAGTGTGCTCGT | 105 |
| *BMP10* | CCTCTGCCAACATCATTAGGAG | TTTTCGGAGCCCATTAAAACTGA | 77 |
| *HAND2* | ACATCGCCTACCTCATGGAC | TGGTTTTCTTGTCGTTGCTG | 162 |
| *mCherry* | CAAGTTGGACATCACCTCCCAC | ACTTGTACAGCTCGTCCATGC | 104 |

**Supplemental Table S2. Sequence of primers used for qPCR**
